## Supplemental Table 1 for "Processing of the Ribosomal Ubiquitin-Like Fusion Protein FUBI-eS30/FAU is Required for 40S Maturation and Depends on USP36"

| **Key Resources Table** | | | | |
| --- | --- | --- | --- | --- |
| **Reagent type (species) or resource** | **Designation** | **Source or reference** | **Identifiers** | **Additional information** |
| Cell line (*Homo sapiens*) | HEK293 Flp-In T-REx | Thermo Fisher Scientific | Cat# R78007  RRID:CVCL_U427 |  |
| Cell line (*Homo sapiens*) | HEK293 Flp-In T-REx FUBI-eS30-StHA | This paper |  | See Materials and methods, *Cell culture, cell lines, and treatments* |
| Cell line (*Homo sapiens*) | HEK293 Flp-In T-REx FUBI(AA)-eS30-StHA | This paper |  | AA: G73,74A  See Materials and methods, *Cell culture, cell lines, and treatments* |
| Cell line (*Homo sapiens*) | HEK293 Flp-In T-REx FUBI(GV)-eS30-StHA | This paper |  | GV: G74V  See Materials and methods, *Cell culture, cell lines, and treatments* |
| Cell line (*Homo sapiens*) | HEK293 Flp-In T-REx HASt-FUBI-eS30 | This paper |  | See Materials and methods, *Cell culture, cell lines, and treatments* |
| Cell line (*Homo sapiens*) | HEK293 Flp-In T-REx HASt-GFP | (Wyler et al., 2011) DOI: 10.1261/rna.2325911 |  |  |
| Cell line (*Homo sapiens*) | HeLa Flp-In T-REx | (Hafner et al., 2014) DOI: 10.1038/ ncomms5397 |  | Obtained from T. Mayer (University of Konstanz) |
| Cell line (*Homo sapiens*) | HeLa Flp-In T-REx FUBI-eS30-StHA | This paper |  | See Materials and methods, *Cell culture, cell lines, and treatments* |
| Cell line (*Homo sapiens*) | HeLa Flp-In T-REx FUBI(AA)-eS30-StHA | This paper |  | AA: G73,74A  See Materials and methods, *Cell culture, cell lines, and treatments* |
| Cell line (*Homo sapiens*) | HeLa Flp-In T-REx FUBI(GV)-eS30-StHA | This paper |  | GV: G74V  See Materials and methods, *Cell culture, cell lines, and treatments* |
| Cell line (*Homo sapiens*) | HeLa FRT/TetR | Other |  | Obtained from M. Beck (EMBL, Heidelberg) |
| Cell line (*Homo sapiens*) | HeLa FRT/TetR EGFP-USP36 | This paper |  | See Materials and methods, *Cell culture, cell lines, and treatments* |
| Cell line (*Homo sapiens*) | HeLa FRT/TetR EGFP-USP36(CA) | This paper |  | CA: C131A  See Materials and methods, *Cell culture, cell lines, and treatments* |
| Cell line (*Homo sapiens*) | HeLa Kyoto | Other | RRID:CVCL_1922 | Obtained from D. Gerlich (IMBA, Vienna) |
| Antibody | anti-AAMP (rabbit polyclonal) | This paper |  | WB (1:800)  See Materials and methods, *Antibodies* |
| Antibody | anti-β-actin (mouse monoclonal) | Santa Cruz Biotechnology | Cat# sc-47778  RRID:AB_626632 | WB (1:1000) |
| Antibody | anti-C21ORF70 (rabbit polyclonal) | (Montellese et al., 2017) DOI: 10.1093/nar/gkx253 |  | WB (1:500) |
| Antibody | anti-EIF1AD | Proteintech | Cat# 20528-1-AP  RRID: AB_10693533 | WB (1:1000) |
| Antibody | anti-ENP1 (rabbit polyclonal) | (Zemp et al., 2009) DOI: 10.1083/jcb.200904048 |  | IF (1:15,000)  WB (1:1000) |
| Antibody | anti-eS1 (RPS3A) (rabbit polyclonal) | (Wyler et al., 2011) DOI: 10.1261/rna.2325911 |  | WB (1:1000) |
| Antibody | anti-eS26 (RPS26) (rabbit polyclonal) | Abcam | Cat# ab104050 RRID: AB_10710999 | WB (1:500) |
| Antibody | anti-FAU (FUBI-eS30) (rabbit polyclonal) | Abcam | Cat# ab135765 | WB (1:500) |
| Antibody | anti-FUBI (rabbit polyclonal) | This paper |  | WB (1:2000)  See Materials and methods, *Antibodies* |
| Antibody | anti-HA (mouse monoclonal) | Enzo Life Sciences | ENZ-ABS120-0200 | IF (1:2000)  WB (1:1000) |
| Antibody | anti-His (mouse monoclonal) | Sigma-Aldrich | Cat# H1029  RRID:AB_260015 | WB (1:2000) |
| Antibody | anti-LSG1 (rabbit polyclonal) | (Wyler et al., 2014) DOI: 10.1016/j.febslet.2014.08.013 |  | WB (1:3000) |
| Antibody | anti-LTV1 (rabbit polyclonal) | (Zemp et al., 2009) DOI: 10.1083/jcb.200904048 |  | IF (1:4000)  IB (1:2000) |
| Antibody | anti-NMD3 (rabbit polyclonal) | (Zemp et al., 2009) DOI: 10.1083/jcb.200904048 |  | IF (1:1000)  IB (1:10,000) |
| Antibody | anti-NOB1 (rabbit polyclonal) | (Zemp et al., 2009) DOI: 10.1083/jcb.200904048 |  | IF (1:5000)  WB (1:2000) |
| Antibody | anti-NOC4L (rabbit polyclonal) | (Wyler et al., 2011) DOI: 10.1261/rna.2325911 |  | WB (1:5000) |
| Antibody | anti-PNO1 (DIM2) (rabbit polyclonal) | (Zemp et al., 2009) DOI: 10.1083/jcb.200904048 |  | IF (1:2000)  WB (1:2000) |
| Antibody | anti-RIOK1 (rabbit polyclonal) | (Widmann et al., 2012) DOI: 10.1091/mbc.E11-07-0639 |  | IF (1:8000)  WB (1:1000) |
| Antibody | anti-RIOK2 (rabbit polyclonal) | (Zemp et al., 2009) DOI: 10.1083/jcb.200904048 |  | IF (1:5000)  WB (1:5000) |
| Antibody | anti-RRP12 (rabbit polyclonal) | (Wyler et al., 2011) DOI: 10.1261/rna.2325911 |  | IF (1:2000)  WB (1:1000) |
| Antibody | anti-Strep (mouse monoclonal) | IBA GmbH | Cat# 2-1507-001  RRID:AB_513133 | WB (1:1000) |
| Antibody | anti-TRIP4 (rabbit polyclonal) | This paper |  | WB (1:50,000)  See Materials and methods, *Antibodies* |
| Antibody | anti-TSR1 (rabbit polyclonal) | (Zemp et al., 2014) DOI: 10.1242/jcs.138719 |  | WB (1:10,000) |
| Antibody | anti-uL23 (RPL23A) (rabbit polyclonal) | (Wyler et al., 2011) DOI: 10.1261/rna.2325911 |  | WB (1:200) |
| Antibody | anti-uS3 (RPS3) (rabbit polyclonal) | (Zemp et al., 2009) DOI: 10.1083/jcb.200904048 |  | WB (1:1000) |
| Antibody | anti-USP10 (rabbit polyclonal) | Sigma-Aldrich | Cat# HPA006731  RRID:AB_1080495 | WB (1:1000) |
| Antibody | anti-USP16 (rabbit polyclonal) | Bethyl Laboratories | Cat# A301-615A  RRID:AB_1211387 | WB (1:500) |
| Antibody | anti-USP36 (rabbit polyclonal) | Sigma-Aldrich | Cat# HPA012082  RRID:AB_1858682 | IF (1:1000)  WB (1:250) |
| Antibody | goat anti-mouse Alexa Fluor 594 | Thermo Fisher Scientific | A-11005, RRID:AB_2534073 | IF (1:250) |
| Antibody | goat anti-rabbit Alexa Fluor 488 | Thermo Fisher Scientific | A-11008, RRID:AB_143165 | IF (1:250) |
| Antibody | goat anti-mouse Alexa Fluor Plus 680 | Thermo Fisher Scientific | Cat# A32729  RRID:AB_2633278 | WB (1:10,000) |
| Antibody | goat anti-rabbit Alexa Fluor Plus 800 | Thermo Fisher Scientific | Cat# A32735, RRID:AB_2633284 | WB (1:10,000) |
| Recombinant DNA reagent | pC2Pi/control | (Boneberg et al., 2019) DOI: 10.1261/rna.069609.118 |  | See Materials and methods, *Molecular cloning* |
| Recombinant DNA reagent | pC2Pi/USP36i-g1 | This paper |  | USP36 guide 1  See Materials and methods, *Molecular cloning* |
| Recombinant DNA reagent | pC2Pi/USP36i-g2 | This paper |  | USP36 guide 2  See Materials and methods, *Molecular cloning* |
| Recombinant DNA reagent | pCDNA5/FRT/TO/FUBI-eS30-StHA | This paper |  | See Materials and methods, *Molecular cloning* |
| Recombinant DNA reagent | pCDNA5/FRT/TO/FUBI(AA)-eS30-StHA | This paper |  | AA: G73,74A  See Materials and methods, *Molecular cloning* |
| Recombinant DNA reagent | pCDNA5/FRT/TO/FUBI(GV)-eS30-StHA | This paper |  | GV: G74V  See Materials and methods, *Molecular cloning* |
| Recombinant DNA reagent | pCDNA5/FRT/TO/EGFP-USP36 | This paper |  | See Materials and methods, *Molecular cloning* |
| Recombinant DNA reagent | pCDNA5/FRT/TO/EGFP-USP36(CA) | This paper |  | CA: C131A  See Materials and methods, *Molecular cloning* |
| Recombinant DNA reagent | pCDNA5/FRT/TO/HASt-FUBI-eS30 | This paper |  | See Materials and methods, *Molecular cloning* |
| Recombinant DNA reagent | pDEST/LTR/Flag-HA-USP36 | Addgene  (Sowa et al., 2009) DOI: 10.1016/j.cell.2009.04.042 | Cat# 22579  RRID:Addgene_22579 | See Materials and methods, *Molecular cloning* |
| Recombinant DNA reagent | pET-28b(+)/His_6_-FUBI-EGFP | This paper |  | See Materials and methods, *Molecular cloning* |
| Recombinant DNA reagent | pET-28b(+)/His_6_-Ub-EGFP | This paper |  | See Materials and methods, *Molecular cloning* |
| Recombinant DNA reagent | pFBD/EGFP/His_10_-USP36-TEV-St | This paper |  | pFBD: pFastBac Dual  See Materials and methods, *Molecular cloning* |
| Recombinant DNA reagent | pFBD/EGFP/His_10_-USP36(CA)-TEV-St | This paper |  | pFBD: pFastBac Dual, CA: C131A  See Materials and methods, *Molecular cloning* |
| Recombinant DNA reagent | pQE30/His_6_-FUBI-eS30 | This paper |  | See Materials and methods, *Molecular cloning* |
| Recombinant DNA reagent | pQE30/His_6_-FUBI(AA)-eS30 | This paper |  | AA: G73,74A  See Materials and methods, *Molecular cloning* |
| Sequence-based reagent | QuikChange primers for FUBI(AA)-eS30 | Sigma-Aldrich |  | 5'-GCAGGCCGCATGCTTGcAGcTAAAGTTCATGGTTCC-3', 5'-GGAACCATGAACTTTAgCTgCAAGCATGCGGCCTGC-3'  See Materials and methods, *Molecular cloning* |
| Sequence-based reagent | QuikChange primers for FUBI(GV)-eS30 | Sigma-Aldrich |  | 5'-GGCCGCATGCTTGGAGtTAAAGTTCATGGTTCC-3', 5'-GGAACCATGAACTTTAaCTCCAAGCATGCGGCC-3'  See Materials and methods, *Molecular cloning* |
| Sequence-based reagent | QuikChange primers for USP36(CA) | Sigma-Aldrich |  | 5'-CCACAACCTaGGCAACACCgcCTTTCTCAATGCCACC-3', 5'-GGTGGCATTGAGAAAGgcGGTGTTGCCtAGGTTGTGG-3'  See Materials and methods, *Molecular cloning* |
| Sequence-based reagent | QuikChange primers for USP36(NdeI) | Sigma-Aldrich |  | 5’- CCGTGTGCAAGAGCGTCagcGAtACaTAtGACCCCTACTTGGAC-3’, 5'-GTCCAAGTAGGGGTCaTAtGTaTCgctGACGCTCTTGCACACGG-3'  See Materials and methods, *Molecular cloning* |
| Sequence-based reagent | 5'ITS1 | Microsynth  (Rouquette et al., 2005) DOI: 10.1038/sj.emboj.7600752 |  | 5'-CCTCGCCCTCCGGGCTCCGTTAATGATC-3'  See Materials and methods, *Northern blot analysis* |
| Sequence-based reagent | ITS2 | Microsynth  (Rouquette et al., 2005) DOI: 10.1038/sj.emboj.7600752 |  | 5'-GCGCGACGGCGGACGACACCGCGGCGTC-3'  See Materials and methods, *Northern blot analysis* |
| Sequence-based reagent | Cy3-5'ITS1 | Microsynth  (Rouquette et al., 2005) DOI: 10.1038/sj.emboj.7600752 |  | 5'-CCTCGCCCTCCGGGCTCCGTTAATGATC-3'  See Materials and methods, *Fluorescence in situ hybridization* |
| Sequence-based reagent | si-AAMP | Microsynth |  | 5’-CAGGAUGGCAGCUUGAUCCUA-3  See Materials and methods, *RNA interference* |
| Sequence-based reagent | si-control | Qiagen | Cat# 1027281 | Allstars Negative Control siRNA  See Materials and methods, *RNA interference* |
| Sequence-based reagent | si-eS4X (RPS4X) | Qiagen |  | 5’-CUGGAGGUGCUAACCUAGGAA-3’  See Materials and methods, *RNA interference* |
| Sequence-based reagent | si-FAU (FUBI-eS30) | Qiagen |  | 5’-CCGGCGCUUUGUCAACGUUGU-3’  See Materials and methods, *RNA interference* |
| Sequence-based reagent | si-uS19 (RPS15) | Microsynth  (Rouquette et al., 2005) DOI: 10.1038/sj.emboj.7600752 |  | 5’-UCACCUACAAGCCCGUAAA-3’  See Materials and methods, *RNA interference* |
| Sequence-based reagent | si-USP10-1 | Qiagen |  | 5'-UCGCUUUGGAUGGAAGUUCUA-3'  See Materials and methods, *RNA interference* |
| Sequence-based reagent | si-USP10-2 | Qiagen |  | 5’-UACGUCAACACCCAUGAUAGA-3’  See Materials and methods, *RNA interference* |
| Sequence-based reagent | si-USP10-3 | Qiagen |  | 5’-AACACAGCUUCUGUUGACUCU-3’  See Materials and methods, *RNA interference* |
| Sequence-based reagent | si-USP10-4 | Qiagen |  | 5’-AAGAACUAGUUCUUACUUCAA-3’  See Materials and methods, *RNA interference* |
| Sequence-based reagent | si-USP36-1 | Qiagen, Microsynth |  | 5’-CAAGAGCGUCUCGGACACCUA-3’  See Materials and methods, *RNA interference* |
| Sequence-based reagent | si-USP36-2 | Qiagen |  | 5’-UCCGUAUAUGUCCCAGAAUAA-3’  See Materials and methods, *RNA interference* |
| Sequence-based reagent | si-USP36-3 | Qiagen |  | 5’-CCGCAUCGAGAUGCCAUGCAU-3’  See Materials and methods, *RNA interference* |
| Sequence-based reagent | si-USP36-4 | Qiagen |  | 5’-UUCCUUGUGAGUAGCUCUCAA-3’  See Materials and methods, *RNA interference* |
| Sequence-based reagent | si-XPO1 (CRM1) | Microsynth  (Zemp et al., 2009) DOI: 10.1083/jcb.200904048 |  | 5’-UGUGGUGAAUUGCUUAUAC-3’  See Materials and methods, *RNA interference* |
| Chemical compound, drug | cycloheximide, CHX | Sigma-Aldrich | Cat# C7698 |  |
| Chemical compound, drug | Leptomycin B | LC Laboratories | Cat# L-6100 |  |
| Chemical compound, drug | tetracycline | Invitrogen | Cat# 550205 |  |

Boneberg, F.M., T. Brandmann, L. Kobel, J. van den Heuvel, K. Bargsten, L. Bammert, U. Kutay, and M. Jinek. 2019. Molecular mechanism of the RNA helicase DHX37 and its activation by UTP14A in ribosome biogenesis. RNA. 25:685-701.

Hafner, J., M.I. Mayr, M.M. Mockel, and T.U. Mayer. 2014. Pre-anaphase chromosome oscillations are regulated by the antagonistic activities of Cdk1 and PP1 on Kif18A. Nat Commun. 5:4397.

Montellese, C., N. Montel-Lehry, A.K. Henras, U. Kutay, P.E. Gleizes, and M.F. O'Donohue. 2017. Poly(A)-specific ribonuclease is a nuclear ribosome biogenesis factor involved in human 18S rRNA maturation. Nucleic Acids Res. 45:6822-6836.

Rouquette, J., V. Choesmel, and P.E. Gleizes. 2005. Nuclear export and cytoplasmic processing of precursors to the 40S ribosomal subunits in mammalian cells. EMBO J. 24:2862-2872.

Sowa, M.E., E.J. Bennett, S.P. Gygi, and J.W. Harper. 2009. Defining the human deubiquitinating enzyme interaction landscape. Cell. 138:389-403.

Widmann, B., F. Wandrey, L. Badertscher, E. Wyler, J. Pfannstiel, I. Zemp, and U. Kutay. 2012. The kinase activity of human Rio1 is required for final steps of cytoplasmic maturation of 40S subunits. Mol Biol Cell. 23:22-35.

Wyler, E., F. Wandrey, L. Badertscher, C. Montellese, D. Alper, and U. Kutay. 2014. The beta-isoform of the BRCA2 and CDKN1A(p21)-interacting protein (BCCIP) stabilizes nuclear RPL23/uL14. FEBS Lett. 588:3685-3691.

Wyler, E., M. Zimmermann, B. Widmann, M. Gstaiger, J. Pfannstiel, U. Kutay, and I. Zemp. 2011. Tandem affinity purification combined with inducible shRNA expression as a tool to study the maturation of macromolecular assemblies. RNA. 17:189-200.

Zemp, I., F. Wandrey, S. Rao, C. Ashiono, E. Wyler, C. Montellese, and U. Kutay. 2014. CK1delta and CK1epsilon are components of human 40S subunit precursors required for cytoplasmic 40S maturation. J Cell Sci. 127:1242-1253.

Zemp, I., T. Wild, M.F. O'Donohue, F. Wandrey, B. Widmann, P.E. Gleizes, and U. Kutay. 2009. Distinct cytoplasmic maturation steps of 40S ribosomal subunit precursors require hRio2. J Cell Biol. 185:1167-1180.
